# The first report and biological characteristics of *Helcococcus ovis* isolated from clinical bovine mastitis in a Chinese dairy herd

**DOI:** 10.1101/2021.04.08.439114

**Authors:** Kai Liu, Zhaoju Deng, Limei Zhang, Xiaolong Gu, Gang Liu, Yang Liu, Peng Chen, Jian Gao, Bo Han, Weijie Qu

## Abstract

*Helcococcus ovis* (*H. ovis*) was first reported in ovine subclinical mastitis milk and post-mortem examinated organs in Spain and the United Kingdom in 1999, subsequently, it appeared in cattle, horse, goat, and human. However, isolation and characterization of the strain in clinical bovine mastitis is unknown. Therefore, our objective was to identify the pathogen in clinical bovine mastitis. A total of 4 strains from 34 bovine mastitis milk samples were identified, there are tiny and transparent colonies from clinical bovine mastitis milk samples in a Chinese dairy farm, however, these colonies could not be identified using on-farm biochemical tests. The isolates were transported to Mastitis Diagnostic Laboratory of China Agricultural University in Beijing. The colonies were identified as a mixture of *H. ovis* and *Arcanobacterium pyogenes* according to microscopic examination and 16S rRNA gene sequencing and the phylogenetic tree was constructed using 16S rRNA gene sequence of *H. ovis* isolates. In addition, the growth curve and biochemistry test were performed, we also examined the antimicrobial resistance profiles and constructed murine mammary infection model. Our results showed that the *H. ovis* were closely related to the strains isolated from China and Japan, growth speed of *H. ovis* was relatively slower than *Strep*.*agalactiae*, the phenotypic characteristics were similar to CCUG37441 and CCUG39041 except to lactose, isolates were sensitive to most of antimicrobials except daptomycin, *H. ovis* could lead to murine mastitis. In this report, we firstly described the characteristics of *H. ovis* that are associated with clinical bovine mastitis in China.

## Introduction

Bovine mastitis is one of the most costly diseases in dairy industry because of the discarding of milk and costs on treatments, as well as culling of cows (1, 2, 3). In China, the most frequently isolated mastitis pathogens were *Escherichia coli* (14.4%), *Klebsiella* spp. (13.0%), coagulase-negative staphylococci (11.3%), *Streptococcus dysgalactiae* (10.5%), and *Staphylococcus aureus* (10.2%) (4). Due to the constraints of on-farm diagnostic tools, some mastitis pathogens that are rarely found in practice could not be accurately identified by on-farm laboratories in China. This might result in false negative results for mastitis diagnosis. *Helcococcus ovis* (***H. ovis***) was first isolated from colonies mixed with *Trueperella pyogenes* (***T. pyogenes***) and *Staphylococcus* spp. from lung, liver, spleen and mastitic milk of two sheep in UK and Spain (5). Subsequently, it was also isolated from cows with abortions (6), puerperal metritis (7, 8), valvular endocarditis (9), horse with pulmonary abscess (10) and sheep with pleuritis and bronchopneumonia (11, 12). Recently, *H. ovis* was found in human with pyogenic disease (13). Schwaiger *et al*. (2012) reported that *H. ovis* together with *Arcanobacterium pyogenes* was detected by PCR, however, they failed to isolate the bacteria from the samples (14).

The objective of this research was to describe the *H. ovis* found in clinical mastitis cases for the first time in China, meanwhile, we determined the phylogeny relation to *H. ovis* isolated from other host species and investigated the antimicrobial resistance profiles.

## Materials and Methods

### Statement of Ethics

All experiments followed the Regulations of Experimental Animals (2008) issued by China Ministry of Science and Technology. All animal procedures were approved by the Institutional Animal Care and Use Committee of Yunnan Agricultural University (Approval No: IACUC-20132030301).

### Isolates of Bacteria and Identification

A total of 34 milk samples with clinical mastitis were collected from a dairy farm in Hebei province and on-farm identification of bacteria were performed on 31^st^ July 2019. In short, single colony of each sample was classified by Gram-staining, then coagulase test was performed on gram-positive *cocci* isolates by using rabbit serum and these isolates were classified as *Staphylococci* spp. and *Streptococci* spp.. Finally, the *Streptococcus* spp. were grouped by Streptococcal grouping kit (Lancefield’s classification kit, Hopebio, Qingdao, China). However, there were four blood agar plates with single tiny and transparent colonies, and were classified into gram positive *cocci* by gram-staining but could not be classified further into *Staphylococcus* spp. nor *Streptococcus* spp.

In order to identify these isolates, the colonies on those four blood agar plates were delivered to Mastitis Diagnostic Laboratory in China Agriculture University (Beijing). The pin-point colonies were transplanted onto blood agar plates and were incubated in 37 °C until 36 h. Heavy growth colonies were observed, some of them were tiny and transparent, while other colonies were white. All the 2 forms of single colonies were exposed to gram-staining and microscope examination. Partial 16S rRNA gene (1300 bp) sequencing was conducted (Forward primer: 5^’^-TACCTTGTTACGACTT-3’, Reverse primer: 5’-AGAGTTTGATCCTGGCTCAG-3’) (15), these sequences were BLAST with the available sequences in Genbank. The phylogenetic tree of these *H. ovis* was constructed using the clustal V method (DNASTAR Lasergene-Megalign, version 7.1).

### Growth Curve of H.ovis

The growth curves of *H. ovis* and *Strep. agalactiae* were assessed simultaneously. For each isolate, 1 mL of bacterial fluid was diluted in 100 mL brain heart infusion broth with 5% fetal bovine serum and placed on a constant temperature shaker (37°C, 220 rpm). 3 mL of bacterial suspension was collected at 0, 2, 4, 6, 8, 10, 12, 14, 16, 18, 20, 22 and 24 h respectively, and optical density (**OD**) of bacterial suspension was determined at 600 nm in a UV spectrophotometer (Jingke Scientific Instrument Co., Ltd., Shanghai, China).

### Biochemistry Test

Biochemical tests were performed on these 4 isolates using HBI biochemical kit according to the manufacturer’s instructions (Hopebio, Qingdao, China).

### Antimicrobial Resistance Test

Antimicrobial resistance tests were performed on these 4 isolates using broth microdilution method and *Streptococcus pneumoniae* ATCC 49619 was used as quality control strain according to Clinical and Laboratory Standards Institute (16). Briefly, plates were incubated at 37°C in a humidified atmosphere for 24 hours. The MIC of each strain was defined as the lowest concentration of an antimicrobial that completely inhibited growth in broth (no growth) after 24 hours of incubation. MIC of penicillin, cefalexin, oxacillin, clindamycin, tetracycline, enrofloxacin, daptomycin, erythromycin and vancomycin against *H. ovis* isolates were determined following CLSI’s guidelines. All antimicrobials agents were ranging concentrations from 0.015 to 16 μg/mL. Isolates were classified following the clinical breakpoints described in CLSI (2019).

### Murine Mammary Infection Model for H.ovis

Pregnant (20 d of gestation) 6–8-wk-old SPF BALB/c mice (SiPeiFu Laboratory Animal Technology, Beijing, China) were used to determine the pathogenic role of *H*.*ovis* during intramammary infection, as described (17). On the third day after parturition, mice were anesthetized with intramuscular injection of 50 mg/kg Zoletil 50 (Virbac, Carros, France). Two groups (n = 15 per group) of mice were allocated with 1 challenge group and 1 negative control group (sterile PBS). Teats ducts of both the L4 (left) and R4 (right) abdominal mammary glands were exposed under a binocular stereoscopic microscope and 100 μL of bacterial suspension (10^5^ CFU/mL) was injected using a syringe with a 34 G blunt needle. Mammary gland tissue was separated and fixed in 5% of paraformaldehyde at 12, 24, and 36 h after challenge (5 mice for each time point). After embedded, sectioned and stained with hematoxylin-eosin, histological evaluation was performed to assess tissue necrosis, polymorphonuclear neutrophilic granulocyte inflammation (i.e., neutrophilic inflammation) and lymphocytic inflammation.

## Results

### Bacteria Isolation and Identification

The small and transparent colonies were gram-positive *cocci* singly or in pairs, while the white colonies were gram-positive singly irregularity rod-shaped bacterium (Figure 1). and those gram-positive *cocci* were identified as *H. ovis*, while the rod-shaped bacterium were identified as *T. pyogenes* (information of 4 isolates was shown in table 1). The sequences of *H*.*ovis* was submitted to Genbank (https://www.ncbi.nlm.nih.gov/genbank/) with the accession number: MT758192.1;MT758194.1; MT758195.1; MT758196.1. All 4 isolates (bold fonts) were closely related to the strains isolated from goat case (*H. ovis*-YYQ1403) and bovine case (*H. ovis*-XJDY-N1-3) in China, meanwhile they were also closely related to strains isolated from swine and bovine cases in Japan (e.g. *H. ovis*-Ymagata-080813, *H. ovis*-Ymagata-160927) (Figure 2).

**Figure 1.**
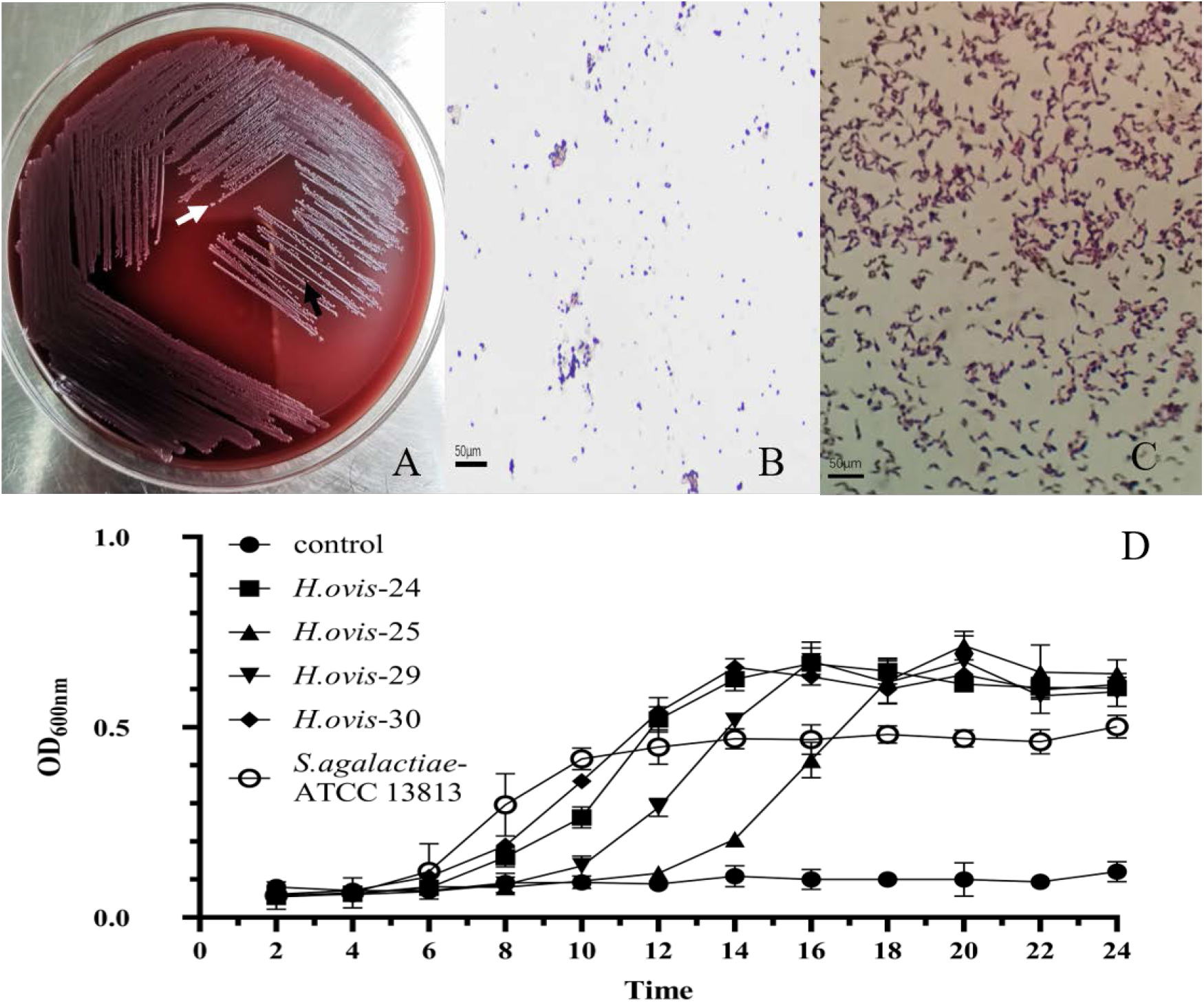
Morphological characteristics and growth curves of *Helcococcus ovis*. (A) White arrow shows the tiny, transparent colony, the black arrow indicates white colony. (B) Gram-positive cocci(400×). (C) Gram-positive irregularity rod-shaped bacterium(400×).

**Table 1.**
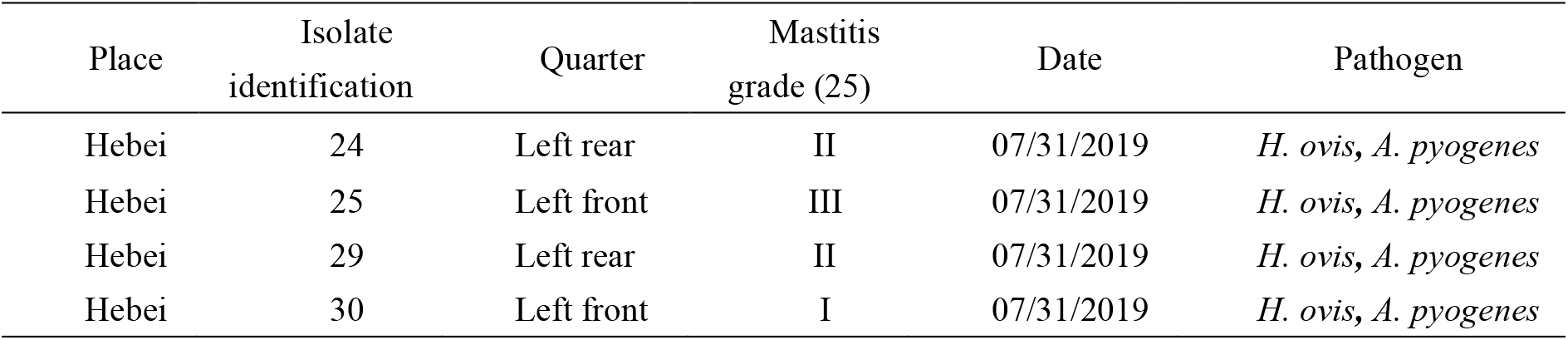
Information of 4 isolates of *H. ovis*.

**Figure 2.**
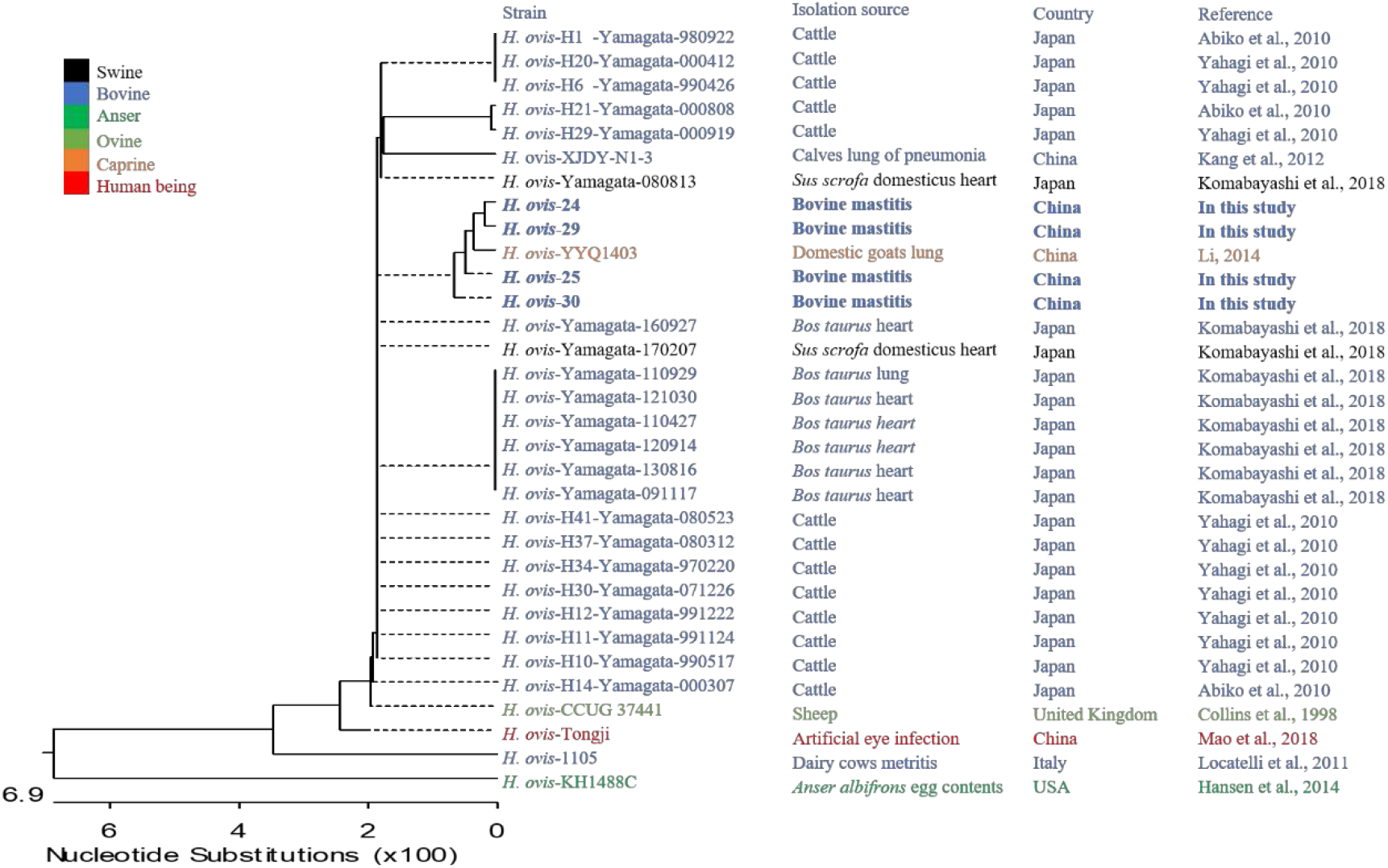
Phylogenetic analysis of 4 strains of *Helcococcus ovis*. Bold indicates *Helcococcus ovis* isolates in this study.

### Growth Curve of H.ovis

The growth curve of *H*.*ovis* has been evaluated with determinations of the OD value. Results suggesting that the bacterial-growth curve of *H. ovis* consisted of a lag phase (∼6 h), a log phase (∼10 h) and a stationary phase. Meanwhile, *H. ovis* had a long lag phase and relatively higher OD_600nm_ value compared to *Streptococcus agalactiae* (Figure 1D).

### Biochemistry Test

Ribose, Mannitol, Sorbitol, Raffinose, Melibiose, Sucrose, β-Mannosidase, Urease, Hippurate, hydrolysis and Esterase reaction were negative except Lactose and Maltose (Table 2). characterization of *H*.*ovis* in this study was similar to the result of *H*.*ovis* CCUG 37441 and CCUG 39041 except to lactose and esterase.

**Table 2.**
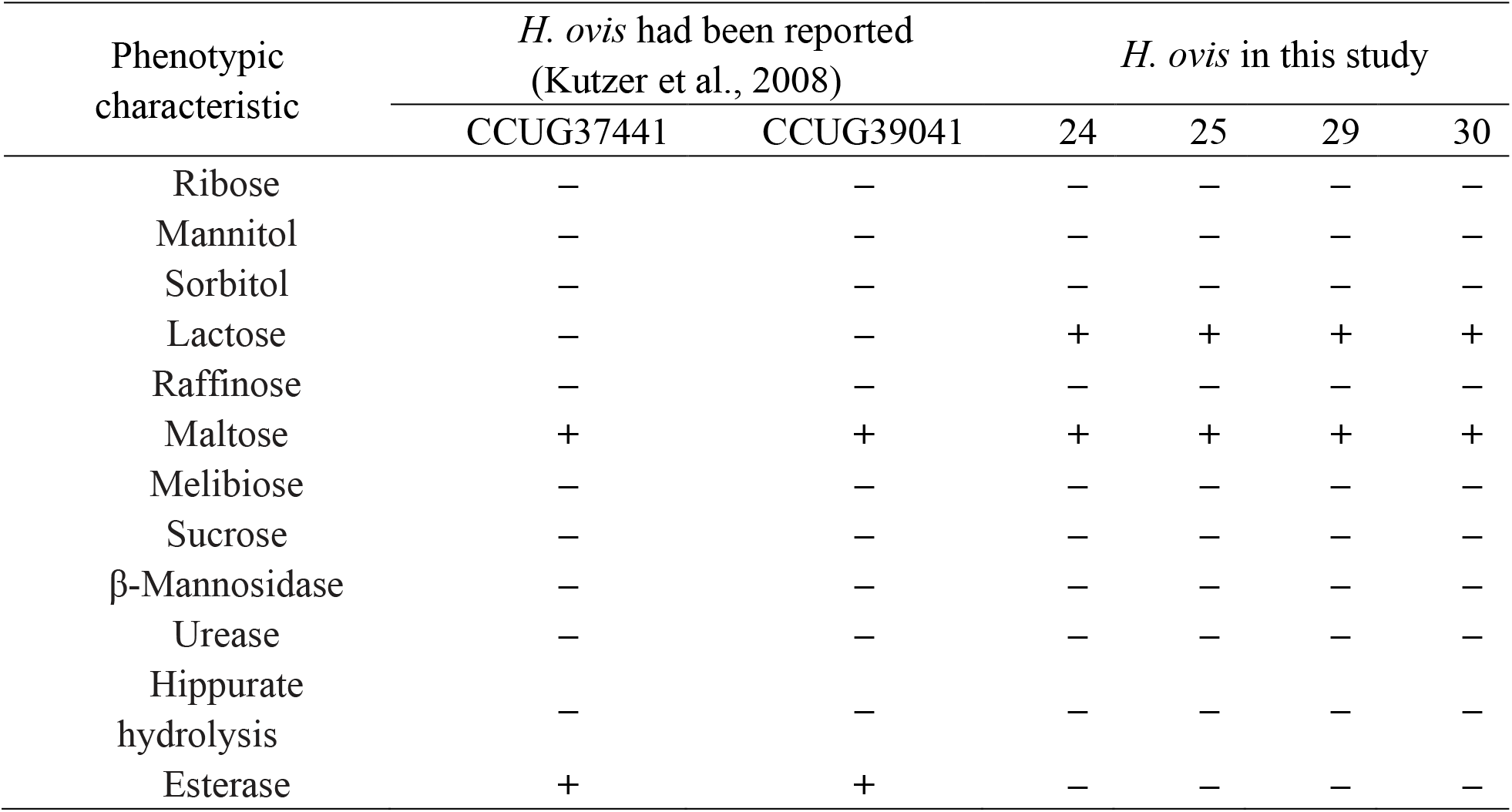
Biochemical results of four *H. ovis* isolates compared to results obtained for *H. ovis* control strains.

### Antimicrobial Resistance Test

In our research, four isolates of *H*.*ovis* were only resistant to daptomycin, and one of isolates were intermediary to tetracycline and enrofloxacin, which were clinically applied to bovine mastitis. As shown in Table 3, results showed that four *H. ovis* isolates exhibiting resistance to daptomycin, while only one isolate (Hebei-24) was intermediate to tetracycline and enrofloxacin. However, all isolates were susceptible to penicillin, cefalexin, oxacillin, clindamycin, erythromycin and vancomycin. One isolate (Hebei-24) was resistant to 3 classes of antimicrobial agents which were defined as multi-drug-resistance (MDR).

**Table 3.** Antimicrobial resistance profiles of the 4 *Helcococcus ovis* isolates.

| Isolates | MIC ($\mu\text{g mL}^{-1}$) | | | | Breakpoints ($\mu\text{g mL}^{-1}$) | | |
| --- | --- | --- | --- | --- | --- | --- | --- |
|  | Hebei-24 | Hebei-25 | Hebei-29 | Hebei-30 | S | I | R |
| Panicillin | 0.015 | 0.015 | 0.015 | 0.015 | $\leq 0.12$ | — | $\geq 0.25$ |
| cefalexin | 1.000 | 1.000 | 1.000 | 0.500 | — | — | $\geq 8$ |
| ceftiofor | $<0.015$ | $<0.015$ | $<0.015$ | $<0.015$ | — | — | $\geq 8$ |
| oxacillin | 0.015 | $<0.015$ | $<0.015$ | $<0.015$ | $\leq 2$ | — | $\geq 4$ |
| clindamycin | 0.250 | 0.250 | 0.250 | 0.250 | $\leq 0.5$ | 1-2 | $\geq 4$ |
| tetracycline | 8.000 | 0.500 | 0.500 | 0.500 | $\leq 4$ | 8 | $\geq 16$ |
| enrofloxacin | 4.000 | 0.500 | 1.000 | 0.500 | $\leq 2$ | 4 | $\geq 8$ |
| daptomycin | $>16.000$ | 16.000 | 16.000 | $>16$ | $\leq 1$ | — | — |
| erythromycin | $<0.015$ | $<0.015$ | $<0.015$ | $<0.015$ | $\leq 0.5$ | 1-4 | $\geq 8$ |
| vancomycin | 0.125 | 0.125 | 0.250 | 0.125 | $\leq 2$ | 4-8 | $\geq 16$ |

Note: S: susceptible; I: intermediate; R: resistance. The breakpoints refers to *Staphylococcus aureus* (CLSI, 2019) were used as (Mao et al., 2018) demonstrated: penicillin $\geq 0.25\mu\text{g mL}^{-1}$ ; cefalexin $\geq 8\mu\text{g mL}^{-1}$ ; oxacillin $\geq 4\mu\text{g mL}^{-1}$ ; clindamycin $\geq 4\mu\text{g mL}^{-1}$ ; tetracycline $\geq 16\mu\text{g mL}^{-1}$ ; enrofloxacin $\geq 8\mu\text{g mL}^{-1}$ ; daptomycin $\geq 1\mu\text{g mL}^{-1}$ ; erythromycin $\geq 8\mu\text{g mL}^{-1}$ and vancomycin $\geq 16\mu\text{g mL}^{-1}$ . Unshaded cells indicate that the MIC value was susceptible, light gray shading indicates intermediate, and green shading indicates resistance.

### Inflammation of Murine Mammary Gland Infected by H.ovis

Swollen and hyperemia mammary gland were observed after 12 h challenged with H.ovis, with more profound pathological changes after 24 h and 36 h (Figure 3A). Histological characteristics of murine mammary glands infected H.ovis were assessed (Figure 3B). Slight infiltrations of inflammatory cells and progressive mammary alveolar damage was observed, stromal hyperplasia appeared in the infected mammary gland after 12 h and 24h challenged with H.ovis, while numerous lymphocytes and formed a tumor-like structure was observed after 36 h, and mammary epithelial cells showed degeneration and necrosis. The bacterial load was 4.6×10^8^ CFU/g after 12h H.ovis challenged, and increased gradually to 6.8×10^8^ CFU/g and 1.8×10^9^ CFU/g after 24 h and 36 h H.ovis challenged, respectively (Figure 3B).

**Figure 3.**
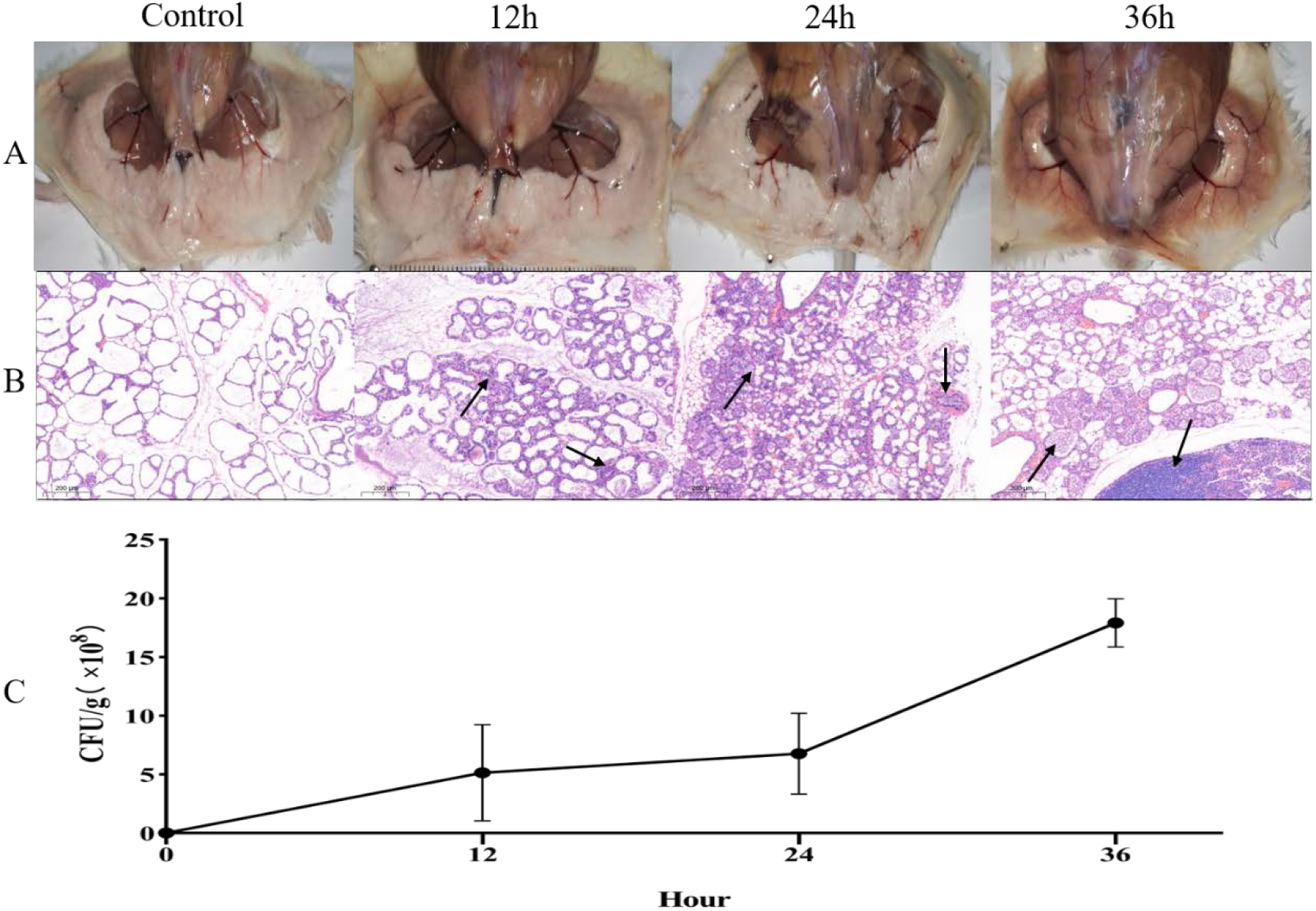
Inflammation and histological evaluation and bacterial burden of murine mammary gland after intramammary inoculation with *Helcococcus ovis*. (A) Pathological changes in murine mammary glands challenged with *Helcococcus ovis*. (B) *Helcococcus ovis* induced murine mastitis, with an increasing number of inflammatory cells (mainly neutrophils) infiltrated into the gland alveoli, stromal hyperplasia and progressive mammary alveolar damage (black arrow) after infection. (C)Bacterial burden in mammary glands of mice (up to 36 h after inoculation). Each time point represents 5 mice in each of the 4 groups. Data are mean ± SD of 3 independent experiments.

## Discussion

To the best of our knowledge, this study is the first report of *H. ovis* isolated from bovine mastitis in China, and is of importance when designing mastitis control plans. In China, there were only four publications of *H. ovis* included three publications of *H. ovis* in goat and bovine pneumonia (18, 19, 12) and one publication in human being artificial eye, which raised the concern of zoonotic property of *H. ovis*(13). Rothschild (2004) assumed this bacteria from skin, whether if it is one the skin microbiota of bovine need further study (10). The colony of *H. ovis* is so tiny that technicians in laboratories may ignore it when it mixes with other major bacteria or milk droplets during routine milk sample processing. Future studies to disclose the pathogenicity of *H. ovis* in causing bovine mastitis are needed.

Bovine mastitis is predominantly caused by bacteria, therefore, antimicrobials is extensively applied to mastitis prevention and treatment (17), which raised the concern of AMR that threatened the health of human beings (20). In our research study, the four isolates of *H*.*ovis* were only resistant to daptomycin, and one of isolates were intermediary to tetracycline and enrofloxacin, which were applied to bovine mastitis. *H*.*ovis* in our study shows slight resistant to antimicrobials, which may indicated that it is rarely exposed to antimicrobial treatment and emerged in mastitis milk shortly. As far as daptomycin, which may attributed to *H*.*ovis* is natural resistant to this drug (21). Phenotypic characteristics of 4 isolates were similar to CCUG 37441 and CCUG 39041 except lactose and esterase (5), this may contribute to the isolates were isolated from mastitis milk and lactose was the main carbohydrate source. Biochemical methods can only be used as an auxiliary inspection method because of its unreliable, molecular identification method can confirm it (13).

The murine model of intramammary challenge with bovine mastitis pathogens has been successfully used to assess bacterial infection and tissue damage (17). The murine model of *H*.*ovis* may improve our research on the correlation between this bacteria and bovine mastitis, the drug effect applied to clinical trials as well as the relationship among bacteria, bovine immune response and lactation (22). In this research, the pathogenicity of *H*.*ovis* on mice at different periods was determined. As time went by, only slight infiltrations of inflammatory cells were observed, stromal hyperplasia appeared, progressive mammary alveolar damage and bacteria burden in tissue appeared which indicated that *H*.*ovis* can not cause severe mastitis. *T*.*pyogenes* was distributed in a lot kinds of animals and can lead to bovine mastitis (23), *H*.*ovis* was isolated together with *T*.*pyogenes*, which may included that *H*.*ovis* may not the major pathogen of bovine mastitis, while can impress the immunity of cows or improve the inflammation (24), further research are needed.

## Acknowledgements

This study was financially supported by the National Natural Science Foundation of China (No. 31660730), Yunnan Expert Workstation (No.202005AF150041).

